# Novel biologically relevant small RNA-sequencing alignment tool *LevenMap* for alignment to database of non-coding RNAs

**DOI:** 10.64898/2026.08.14.742100

**Authors:** Hunter Dlugas, Gregory Dyson, Alan Dombkowski, Yeonju Kim, Katherine Gurdziel, Julie L Boerner, Cathryn Bock

## Abstract

A crucial aspect of the bioinformatics workflow in small RNA-sequencing is the alignment of reads to a database of reference ncRNAs. Alignment algorithms such as Bowtie, Burrows-Wheeler Aligner (BWA), and Spliced Transcripts Alignment to a Reference (STAR) - which are designed for aligning reads to a reference genome - are typically used. Aligning short RNA-sequenced reads to a database of non-coding RNAs (ncRNAs) is fundamentally a different task than aligning longer reads to a genome due to ncRNAs (i) having roughly the same number of nucleotides as the reads being aligned and (ii) being subsequences of other ncRNAs. To account for these differences, we developed the novel alignment algorithm *LevenMap*. Of all reads which exactly matched a reference ncRNA in a publicly available dataset, LevenMap aligned 100.0% of them to their respective ncRNA while all other aligners mapped less than 40% of these reads to their corresponding ncRNA. Furthermore, the mean ratio (length of read) / (length of corresponding reference ncRNA) of all aligned reads was 1.0 and 0.998 for LevenMap with at most zero and one mismatch(es) allowed, respectively; this ratio was no more than 0.51 for all other aligners. Overall, LevenMap is designed to account for the nuances of aligning small RNA-sequencing data to a database of reference ncRNAs and yields more biologically relevant counts compared to traditional aligners in this context. LevenMap is free and publicly available on GitHub: https://github.com/hdlugas/LevenMap.

## 1 Introduction

In recent years, it has become more well-established that non-coding RNAs (ncRNAs) are involved in many aspects of regulating gene expression. Small RNA-sequencing can be used to discover and/or quantify clinically relevant ncRNA biomarkers. In particular, ncRNAs have been discovered and proposed to be biomarkers of various cancers [1], [2], [3], [4], [5], [6], [7], [8], [9], [10], [11].

Common bioinformatics workflows which process small RNA-sequencing data typically involve an alignment algorithm such as Bowtie [12], [13] or Burrows-Wheeler Aligner (BWA) [14] to align reads to a database of ncRNAs such as miRbase [15]. Basic Local Alignment Search Tool (BLAST) [16] and Spliced Transcripts Alignment to a Reference (STAR) [17], while perhaps uncommon in the context of small RNA-sequencing, are also two aligners sometimes used in genomics.

In a database of ncRNAs, it is often the case that one ncRNA contains a subsequence equivalent to a smaller ncRNA. For example, due to mature microRNA (miRNA) being derived from precursor hairpin miRNA, a mature miRNA is typically a subsequence of a hairpin miRNA (e.g. the hairpin miRNA hsa-mir-12117 contains the mature miRNA hsa-miR-12117). If a small RNA-sequenced read is exactly equivalent to a mature miRNA such as hsa-miR-12117 and either Bowtie, BLAST, BWA, or STAR is used to align the given read to a database containing both the mature and hairpin miRNA, then the read will often either be (i) unaligned or (ii) aligned to the hairpin miRNA even though the read exactly matches the mature miRNA. This occurs because these commonly used aligners are designed to align reads to a genome where reference sequences are:

- many orders of magnitude longer than the reads being aligned. For example, sequencing techniques do not produce reads corresponding to entire chromosomes, whereas small RNA-sequencing often does produce reads on the same scale as reference ncRNAs with respect to length.
- not exact subsequences of each other. For example, chromosome 13 is not an exact subsequence of chromosome 17, whereas some reference ncRNAs are exact subsequences of other reference ncRNAs (e.g. mature miRNAs are often exact subsequences of hairpin miRNAs).

For example, in a given sample, suppose there are:

- 1,000 reads with the same nucleotide sequence AAACAAGGGGTTCCATCGG.
- a reference database with:
  – ref-1 = **AAACAAGGGGTTCCATCGG**
  – ref-2 = T**AAACAAGGGGTTCCATCGG**
  – ref-3 = **AAACAAGGGGTTCCATCGG**G
  – ref-4 = TC**AAACAAGGGGTTCCATCGG**CCG
  – ref-5 = TTTTCCCCCA**AAACAAGGGGTTCCATCGG**GGTTTCGGAACATGGGCTCAG… …
GACAGCGGGTGTCA.

Note that the subsequences of the reference RNAs which match the nucleotide sequence of the 1000 reads are bolden and colored red. The alignment of these 1000 duplicate reads to this reference database is summarized in Table 1. Observe that BLAST, Bowtie, BWA, and STAR are not ideal because reads which are an exact match to a reference RNA did not consistently align to the corresponding reference RNA; rather, these reads are either unaligned or appear to be randomly aligned to any one of the five candidate reference RNAs depending on the aligner. On the other hand, LevenMap does align all reads to the reference RNA they exactly match regardless of the number of mismatches allowed. The toy data and a Bash script that reproduces this example can be found on GitHub: https://github.com/hdlugas/LevenMap/tree/main/toy_example_from_manuscript_introduction.

**Table 1:** Toy example alignment of 1000 duplicate reads to five reference sequences, one of which exactly matches the reads (ref-RNA-1), and the other four of which contain the read as an exact proper subsequence. The aligners LevenMap, Bowtie, BWA, STAR, and BLAST are compared. Flag: indicates whether or not to suppress alignments if multiple alignments exist (T: only alignments of reads with exactly one alignment are reported; F: all alignments are reported). L: seed length.

| Aligner | ref-RNA-1 | ref-RNA-2 | ref-RNA-3 | ref-RNA-4 | ref-RNA-5 |
| --- | --- | --- | --- | --- | --- |
| LevenMap:<br>no mismatches | 1000 | 0 | 0 | 0 | 0 |
| LevenMap:<br>at most one mismatch | 1000 | 0 | 0 | 0 | 0 |
| Bowtie:<br>Flag=T, L=28 | 0 | 0 | 0 | 0 | 0 |
| Bowtie:<br>Flag=T, L=10 | 0 | 0 | 0 | 0 | 0 |
| Bowtie:<br>Flag=F, L=28 | 189 | 211 | 202 | 199 | 199 |
| Bowtie:<br>Flag=F, L=10 | 189 | 211 | 202 | 199 | 199 |
| BWA:<br>L=32 | 197 | 209 | 192 | 199 | 203 |
| BWA:<br>L=10 | 197 | 209 | 192 | 199 | 203 |
| STAR | 0 | 0 | 0 | 1000 | 0 |
| BLAST | 0 | 0 | 0 | 0 | 0 |

Many bioinformatic workflows have been proposed for processing small RNA-sequencing data [18], and many of these workflows use an aligner designed for aligning to a genome, not to a database of ncRNAs. For example, CAP-miRseq uses Bowtie as its aligner [19], Potla et al propose using either Bowtie or BWA [20], miRge uses Bowtie, DeAnniso uses BLAST [21], and ShortStack uses Bowtie [22]. The cloud-based tool quagmiR utilizes the Levenshtein distance in three regions of a read (3’ end, 5’ end, and motif) to detect miRNA isomers and requires user-specified motif sequences for each reference miRNA [23]. To circumvent issues related to alignment to a database of ncRNAs with Bowtie, BWA, STAR, and BLAST, we present the novel alignment tool LevenMap. This tool considers the nuances of (i) nested reference sequences and (ii) reference sequences which have approximately the same number of nucleotides as the reads produced from small RNA-sequencing. As its name implies, LevenMap aligns a given read to the reference sequence(s) with minimal Levenshtein distance with respect to the given read (given that this minimal Levenshtein distance is below some user-defined threshold).

## 2 Methods

### 2.1 Algorithm

A flowchart depicting how LevenMap integrates in a common small RNA-seq workflow is shown in Figure 1 (a).

**Figure 1:**
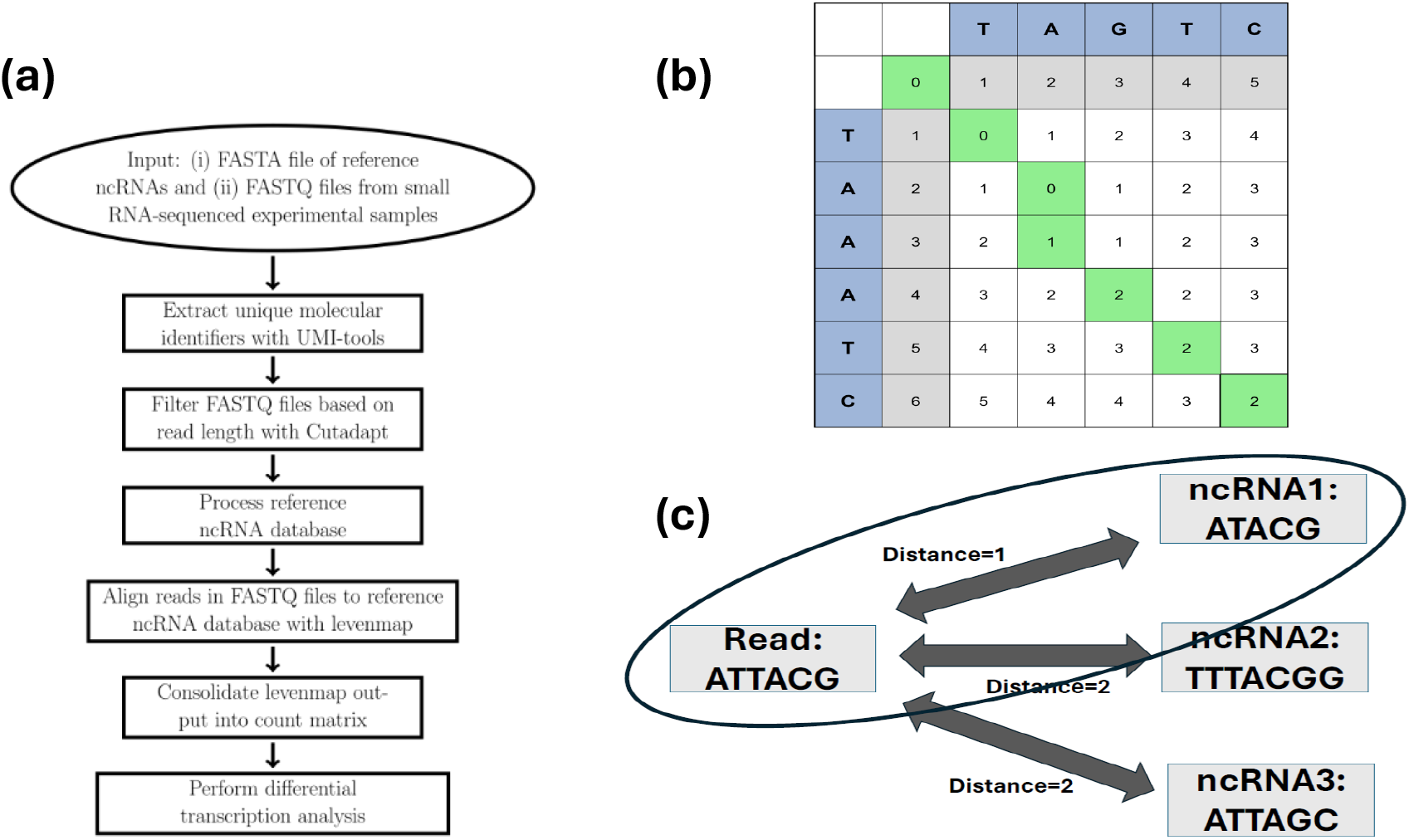
**(a)** Overall small RNA-seq workflow incorporating LevenMap alignment. **(b)** Toy example Levenshtein distance matrix with (*i, j*) entry being the Levenshtein distance between the first *i* characters of the vertical string ‘TAAATC’ and the first *j* characters of the horizontal string ‘TAGTC’. The two strings have a Levenshtein distance of 2, and the green path represents one such way to see this: insert ‘A’ between ‘A’ and ‘G’ in ‘TAGTC’, then substitute ‘G’ for ‘A’. **(c)** Illustration of how LevenMap aligns a given read to the ncRNA(s) with minimal Levenshtein distance (provided such distance is below user-defined threshold).

#### 2.1.1 Build reference database

Similar to how commonly used aligners such as BLAST, Bowtie, BWA, or STAR require an index of the reference database to be constructed prior to alignment, LevenMap also requires the user to process the reference database prior to alignment. Since LevenMap aligns reads to a reference ncRNA database while allowing for at most N mismatches, only reference ncRNAs of length between L-N and L+N, inclusive, need be considered when aligning a given read of length L to the reference ncRNAs. With m, M denoting the number of nucleotides in the shortest and longest reference ncRNA, respectively, computing (N+1)(M-m+1) FASTA files, each of which contains the reference ncRNAs with length in [*m*^⋆^ *n*^⋆^, *m*^⋆^ + *n*^⋆^] for *m*^⋆^ ∈ { *m, m* + 1, …, *M* − 1, *M*}, *n*^⋆^ ∈ {0, 1, …, *N*}, allows for LevenMap to only consider the relevant ncRNAs for a given read and maximum number of mismatches allowed during alignment. Pseudocode explicitly describing how the reference database is processed prior to computing alignments with LevenMap is given in the Supplementary File.

#### 2.1.2 Alignment

Formally, given a finite alphabet *A* (in our case, *A* = {A,T,C,G}), the Levenshtein distance between two words (finite sequences of characters from *A*) *x* and *y* is *d*(*x, y*) = min{*n* + *m* | *x* can be translated into *y* by *n* insertions or deletions and *m* replacements}.

Intuitively, the Levenshtein distance between *x* and *y* is the number of insertions, deletions, or substitutions required to transform *x* into *y*. As its name implies, LevenMap aligns a particular read - and its reverse complement - to reference ncRNAs with minimal Levenshtein distance given that this minimal Levenshtein distance is below some user-defined threshold N. In doing this, the alignment(s) with the fewest mismatches is (are) guaranteed to be reported if alignments with Levenshtein distance below N exist. For example, if a user inputs N=2 and if a given read is an exact match to reference ncRNA X and is only one mismatch away from reference ncRNA Y, then the read aligns only to ncRNA X. This alignment process is described in pseudocode in the Supplementary File.

After running LevenMap on all samples in some experiment, a count matrix with rows corresponding to ncRNAs and columns corresponding to samples is constructed. Differential expression analysis can then be performed on this count matrix to identify ncRNAs with mean expression level significantly different among groups of samples (e.g. cancer vs non-cancer samples, tumor tissue vs non-tumor tissue samples, etc.). DESeq2, edgeR, and limma are common tools for such a differential expression analysis [24], [25], [26].

### 2.2 Application to real-world data

#### 2.2.1 Datasets

For validation, we contrasted LevenMap to BLAST, Bowtie, BWA, and STAR using two real-world datasets: a publicly available dataset (https://www.ebi.ac.uk/ena/browser/view/SRP335559, hence-forth termed SRP335559) originally curated to investigate the effects of TENT2, TUT4, and TUT7 on miRNA regulation [27] and an in-house prostate cancer cohort. The SRP335559 dataset consists of 12 samples: two TUT4 knockout (KO), two TUT7 KO, two TENT2 KO, two double TUT4/TUT7 KO, two triple TENT2/TUT4/TUT7 KO, and two wild-type (WT). Further details on this database can be found in Yang et al [27].

The prostate cancer dataset consists of 34 men with PI-RADS score ≥3 scheduled for an MRI-guided biopsy at the Karmanos Cancer Center. All patients signed informed consent to be a part of this study (Wayne State University IRB #110117M1F) with methods carried out in accordance with the relevant guidelines and regulations. Characteristics of this cohort are described in Table S1 in the Supplementary File.

A pre-biopsy serum sample was gathered from each of the 34 patients. During MRI-guided biopsy, one sample was obtained from the target lesion with the highest PI-RADS score after a sample was obtained for pathology staging. If there were multiple lesions with the same PI-RADS score, then the largest lesion was targeted. There were 31 targeted site samples gathered in this manner. Additionally, 30 non-targeted site samples were obtained from a site that was both away from the target lesion and that appeared cancer-free on both MRI and ultrasound.

The serum RNA isolation was performed by using the Plasma/Serum RNA Purification maxi kit (Norgen Biotek Corp., Ontario Canada; cat# 56200). DNA removal was included using the RNase-Free DNase I Kit (Norgen; cat# 25710) following the manufacturer’s instructions. The RNA quality and concentration was assessed by using the RNA 6000 Pico Kit (Agilent Technologies, Waldbronn, Germany; cat# 5067-1513) and the 2100 Bioanalyzer (Agilent Technologies).

RNA templates were normalized at 1 or 10ng. Then, NGS library preparation was performed using the QIAseq miRNA kit (QIAGEN, Maryland USA; cat# 1103679) with QIAseq miRNA Index kit UDI-C Version 2 (QIAGEN; cat# 331895) for dual indexing following the manufacturer’s instructions. The libraries were prepared on a Vapo-protect Pro S (Eppendorf, Hamburg, Germany). The high sensitivity D1000 kit (Agilent Technologies; cat# 5067-5584 & 5067-5585) and the 4200 TapeStation, (Agilent Technologies, Waldbronn, Germany) were used to determine library quality, and the concentrations were determined using the Qubit™ dsDNA HS Assay Kit (Life Technologies Corporation, Oregon, USA; cat# Q33231) and Qubit 2.0 (Life Technologies). Libraries were sequenced on the NovaSeq 6000 (72 bp read length).

#### 2.2.2. Bioinformatic approach

The unique molecular identifier (UMI) of each read was extracted using UMI-tools [28]. Cutadapt was then used to remove reads with less than 16 nucleotides or greater than 1559 nucleotides [29]. The thresholds of 16 and 1559 nucleotides were liberally chosen based on the shortest and longest ncRNA in the reference ncRNA database being 16 and 1559 nucleotides, respectively. This reference database consists of both (i) all human hairpin and mature miRNAs in miRbase [15] and (ii) all ncRNAs in tsRNAsearch’s ncRNA database [30]. Overall, this pre-alignment preprocessing is similar to that proposed by Potla et al [20].

Once UMIs were extracted and reads filtered based on length, LevenMap was used to align reads to the reference ncRNA database while allowing for at most one mismatch. For the sake of comparing alignment characteristics, the case of no mismatches was considered as well. Bowtie (v1.3.1) and BWA (v0.7.17) were also used to align reads to the same reference ncRNA database with several parameter combinations similar to recommended parameters for aligning short reads [12], [13], [14]. Additionally, BLAST (v2.10.1) and STAR (v2.7.11) were implemented as well with default parameters [16], [17]. Alignment characteristics among LevenMap, BLAST, Bowtie, BWA, and STAR are then compared.

To illustrate the downstream effect of alignment on biomarker discovery, a differential expression analysis was performed to identify ncRNAs differentially expressed with respect to:

- SRP335559:
  – TENT2 KO vs WT
  – TUT4 KO vs WT
  – TUT7 KO vs WT
  – TUT4/TUT7 KO vs WT
  – TENT2/TUT4/TUT7 KO vs WT
- Prostate cancer cohort:
  – Targeted site cancer status.

To this end, only ncRNAs with a count (i) of at least three in at least two samples and (ii) standard deviation at least three were considered high-quality and retained. The Bioconductor package DESeq2 was then used to normalize the count data using the median of ratios methods [24]. For the SRP335559 dataset, DESeq2 was used to perform negative binomial regression whereas in the prostate cancer cohort, generalized least squares with the within-group correlation structure of targeted site samples and non-targeted site samples taken to be compound symmetry structure corresponding to a constant correlation was used for the differential expression analysis [31], [32]. Differentially expressed ncRNAs were defined as ncRNAs with (i) nominal p-value less than 0.05 and (ii) | log_2_ (fold-change) | larger than log_2_ (2) = 1.

## 3 Results

When at most one mismatch is allowed with LevenMap, sometimes a given read will align to multiple reference ncRNAs if no reference ncRNAs exactly match the read. For example, the read AAAAG-TAATTGCGGTTTTTGCT aligns to both hsa-miR-548am-5p (AAAAGTAATTGCGGTTTTTGC**C**) and hsa-miR-548au-5p (AAAAGTAATTGCGGTTTTTGC) when at most one mismatch is allowed. Across all samples, the number of alignments of unique aligned reads (i.e. all aligned reads with duplicates removed) is summarized in Table 2. As observed in Table 2, roughtly 95% of unique aligned reads have exactly one optimal alignment regardless of dataset.

**Table 2:** Number of optimal alignments of unique reads across all samples when LevenMap allows for at most 1 mismatch. For example, there are 541 unique reads (i.e., not counting duplicates) across all samples in the SRP335559 dataset which aligned to exactly 2 reference ncRNAs.

| Number of alignments | Number of unique reads |  |
| --- | --- | --- |
|  | SRP335559 | Prostate Cancer |
| 1 | 12,503 (95.52%) | 2,752 (94.21%) |
| 2 | 541 (4.13%) | 165 (5.65%) |
| 3 | 38 (0.29%) | 4 (0.14%) |
| 4 | 4 (0.03%) | 0 (0.0%) |
| 5 | 2 (0.02%) | 0 (0.0%) |
| 6 | 1 (0.01%) | 0 (0.0%) |

To demonstrate the improved alignment characteristics of LevenMap compared to BLAST, Bowtie, BWA, and STAR, several statistics are considered:

- ratio of read length to length of corresponding ncRNA for aligned reads
- proportion of aligned reads
- proportion of reads with an exact match which actually align to the exact match
- proportion of reads with an exact match which align to reference ncRNAs containing their exact match as a proper subsequence
- proportion of reads with exact match which align to any reference ncRNA

These alignment characteristics are summarized in Table 3. In the Venn diagrams in Figure S1 of the Supplementary File, observe that there is little to no overlap in the differentially expressed ncRNAs among all aligners, including the case of LevenMap vs the others.

**Table 3:** Summary of alignment characteristics for LevenMap, Bowtie, BWA, STAR, and BLAST. Cells are mean *±* standard deviation across all samples. Ratio = mean 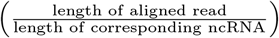 Proportion of aligned reads = mean 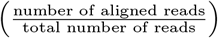 Proportion of reads with exact match which align to the exact match: Where 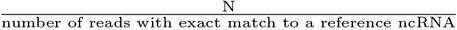 is the number of reads with an exact match to a reference ncRNA which aligned to the matching reference ncRNA. Proportion of reads with exact match which align to ncRNA containing exact match as proper subsequence: 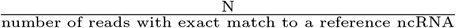 where *N* is the number of reads with an exact match to a reference ncRNA which aligned to a reference ncRNA containing the true reference ncRNA as a proper subsequence. Proportion of reads with exact match which align to any ncRNA: 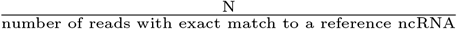 where *N* is the number of reads with exact match to a reference ncRNA which align to any ncRNA. Flag: indicates whether or not to suppress alignments if multiple alignments exist (T: only alignments of reads with exactly one alignment are reported; F: all alignments are reported). L: seed length.

| Dataset | Aligner | Ratio | Proportion of aligned reads | Proportion of reads with exact match which align to the exact match | Proportion of reads with exact match which align to ncRNA containing exact match as proper subsequence | Proportion of reads with exact match which align to any ncRNA |
| --- | --- | --- | --- | --- | --- | --- |
| SRP335559 | LevenMap:<br>no mismatches | 100.0% ± 0.0% | 14.7% ± 9.9% | 100.0% ± 0.0% | 0.0% ± 0.0% | 100.0% ± 0.0% |
| | LevenMap:<br>at most one mismatch | 99.8% ± 0.8% | 27.4% ± 19.0% | 100.0% ± 0.0% | 0.0% ± 0.0% | 100.0% ± 0.0% |
| | LevenMap:<br>at most two mismatches | 99.7% ± 1.1% | 30.9% ± 21.0% | 100.0% ± 0.0% | 0.0% ± 0.0% | 100.0% ± 0.0% |
| | Bowtie:<br>Flag=T, L=28 | 25.1% ± 1.9% | 4.7% ± 3.3% | 0.0% ± 0.0% | 0.0% ± 0.0% | 0.0% ± 0.0% |
| | Bowtie:<br>Flag=T, L=10 | 21.5% ± 3.0% | 6.9% ± 5.2% | 0.0% ± 0.0% | 0.0% ± 0.0% | 0.0% ± 0.0% |
| | Bowtie:<br>Flag=F, L=28 | 41.7% ± 4.7% | 35.4% ± 17.7% | 39.3% ± 3.1% | 58.7% ± 1.4% | 100.0% ± 0.0% |
| | Bowtie:<br>Flag=F, L=10 | 37.9% ± 4.7% | 44.9% ± 23.1% | 39.3% ± 3.1% | 58.7% ± 1.4% | 100.0% ± 0.0% |
| | BWA:<br>L=32 | 35.4% ± 6.5% | 48.4% ± 19.1% | 38.8% ± 3.6% | 56.9% ± 1.5% | 96.6% ± 3.5% |
| | BWA:<br>L=10 | 35.4% ± 6.5% | 48.4% ± 19.1% | 38.8% ± 3.6% | 56.9% ± 1.5% | 96.6% ± 3.5% |
| | STAR | 21.7% ± 1.7% | 58.9% ± 17.6% | 0.0% ± 0.0% | 98.6% ± 2.0% | 98.6% ± 2.0% |
| | BLAST | 26.3% ± 1.7% | 6.1% ± 3.6% | 0.0% ± 0.0% | 0.0% ± 0.0% | 0.0% ± 0.0% |
| Prostate Cancer | LevenMap:<br>no mismatches | 100.0% ± 0.0% | 53.2% ± 4.7% | 100.0% ± 0.0% | 0.0% ± 0.0% | 100.0% ± 0.0% |
| | LevenMap:<br>at most one mismatch | 99.9% ± 0.1% | 77.2% ± 5.8% | 100.0% ± 0.0% | 0.0% ± 0.0% | 100.0% ± 0.0% |
| | LevenMap:<br>at most two mismatches | 100.1% ± 0.2% | 86.5% ± 6.3% | 100.0% ± 0.0% | 0.0% ± 0.0% | 100.0% ± 0.0% |
| | Bowtie:<br>Flag=T, L=28 | 26.6% ± 0.4% | 9.2% ± 1.1% | 0.0% ± 0.0% | 0.0% ± 0.0% | 0.0% ± 0.0% |
| | Bowtie:<br>Flag=T, L=10 | 25.1% ± 0.3% | 8.2% ± 1.5% | 0.0% ± 0.0% | 0.0% ± 0.0% | 0.0% ± 0.0% |
| | Bowtie:<br>Flag=F, L=28 | 50.7% ± 0.7% | 79.8% ± 3.8% | 36.9% ± 1.2% | 59.4% ± 0.8% | 100.0% ± 0.0% |
| | Bowtie:<br>Flag=F, L=10 | 48.9% ± 0.7% | 89.1% ± 4.4% | 36.9% ± 1.2% | 59.4% ± 0.8% | 100.0% ± 0.0% |
| | BWA:<br>L=32 | 50.1% ± 0.9% | 89.0% ± 3.7% | 36.9% ± 1.3% | 55.8% ± 0.6% | 94.8% ± 0.9% |
| | BWA:<br>L=10 | 50.1% ± 0.9% | 89.0% ± 3.7% | 36.9% ± 1.3% | 55.8% ± 0.6% | 94.8% ± 0.9% |
| | STAR | 25.6% ± 0.3% | 93.8% ± 4.2% | 0.0% ± 0.0% | 99.8% ± 0.1% | 99.8% ± 0.1% |
| | BLAST | 16.8% ± 1.2% | 2.0% ± 1.8% | 0.0% ± 0.0% | 0.0% ± 0.0% | 17.1% ± 19.1% |

As observed in Table S2 of the Supplementary File, the computational expense of LevenMap is comparable to STAR while BWA and Bowtie are the fastest and BLAST is the most demanding. With four CPUs, the mean ± standard deviation time required for LevenMap to align each sample while allowing for at most one mismatch is 6.21 ± 2.41 minutes and 16.81 ± 13.11 minutes for the prostate cancer and SRP335559 dataset, respectively. Expectedly, these times are roughly 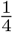 of the respective times when one CPU is used.

The mean 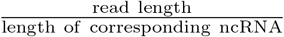 for all aligned reads is significantly different between LevenMap and with each of the four other aligners from two-sample paired t-tests with Welch approximation for all cases: LevenMap vs Bowtie, LevenMap vs BWA, LevenMap vs STAR, and LevenMap vs BLAST (*p <* 0.001). Additionally, for all non-LevenMap aligners, we reject the null hypotheses of the mean and median proportion of reads with an exact match which aligned to the true match being equal to 1 using one-sample t-tests and Wilcoxon signed-rank tests, respectively (*p <* 0.001). Similarly, the null hypotheses of the mean and median proportion of reads with an exact match which aligned to a ncRNA containing the true match as an exact subsequence being equal to 0 can be rejected per one-sample t-tests and Wilcoxon signed-rank tests, respectively (*p <* 0.001).

Although not the primary focus of this work, some differential expression results are summarized in the Supplementary File in Figures S1 (overlap of differentially expressed ncRNAs among aligners), S2 (comparison of ncRNAs significant in LevenMap only and Bowtie only for a single analysis) and S3 (volcano plot of significant results using the LevenMap aligner with at most one mismatch for all analyses).

## 4 Discussion

It is observed from the Venn diagrams in Figure S1 in the Supplementary File depicting the commonality of differentially expressed ncRNAs that there is little agreement of these downstream results among aligners. The heatmaps in Figure S2 in the Supplementary File can help shed some insight into this lack of discrepancy between LevenMap and Bowtie specifically. The ncRNAs differentially expressed with respect to LevenMap but not Bowtie tend to have diluted Bowtie counts due to the true ncRNA detected by LevenMap aligning to the true ncRNA by Bowtie only about 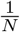 times where *N* is the number of ncRNAs containing the true ncRNA (not necessarily as a proper subsequence). Conversely, there are often non-miRNA ncRNAs that tend to be much longer than miRNAs that are differentially expressed with respect to Bowtie but not LevenMap. This pattern was consistent for all of the alignments compared; TUT4/TUT7 KO vs WT was used as an illustration. Figure S3 shows that the LevenMap alignment results in an increase in the number of significant markers as the number of perturbations increases in the SRP335559 dataset.

If we consider a read with 21 nucleotides and assume (i) a base error rate of 0.024 (per Yang et al [27]) and (ii) that probabilities of bases being called incorrectly are independent, then the probability of the read having no mismatches is 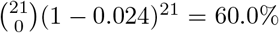 while the probability of the read having at most one mismatch is 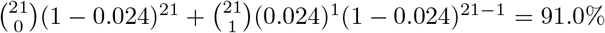 Allowing for at most two mismatches bumps this estimate up by an additive factor of 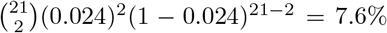 meaning that the probability of the read having at most two mismatches is larger than 98%. With these crude estimates in mind, we recommend setting LevenMap to allow for at most one mismatch as this strikes a fair balance between (i) number of reads aligned and (ii) alignment quality.

The novel LevenMap tool is a more biologically relevant small RNA-sequencing aligner than Bowtie, BWA, STAR, and BLAST in the sense that it aligns reads to their most similar reference ncRNA (if such alignments exist). For example, many reads that align to a reference ncRNA with Bowtie, BWA, and/or STAR align to a reference ncRNA that is more than twice as long as the reads themselves. On the other hand, LevenMap aligns a read to reference ncRNAs within ± *N* nucleotide(s) from the read’s length given that at most *N* mismatch(s) is/are allowed during alignment. As observed in Table 3, BWA, Bowtie without suppressing any alignments, and STAR actually aligned the majority of reads with an exact match to a reference ncRNA that contains the exact match as a proper subsequence rather than to the exact match itself. Additionally, of all reads which have an exact match, LevenMap aligns 100% of them to their true match while BWA, Bowtie, STAR, and BLAST align no more than 40% to their true match on average. Given that (i) most reference ncRNAs which are also contained in a longer reference ncRNA are contained in no more than 2 longer reference ncRNAs and (ii) our observation in the Introduction section that, with Bowtie and BWA, N duplicate reads whose nucleotide sequence is contained in M reference ncRNAs align to each of the M reference ncRNAs roughly 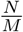 times, the proportion of reads with an exact match which align to their exact match for BWA and Bowtie with all alignments reported being less than 50% may not be surprising.

It is noteworthy to mention that when no mismatches are allowed with LevenMap, there is a smaller mean proportion of reads that align compared to BWA as indicated in Table 3. However, when at most 1 mismatch is allowed with LevenMap, the distribution of the mean proportion of aligned reads across samples is more similar to that of Bowtie (when reads with multiple alignments aren’t suppressed), BWA, and STAR although still slightly smaller. However, given that BLAST, Bowtie, BWA, and STAR do not consistently produce the most biologically relevant alignments, the reduced number of aligned reads with LevenMap may be a not necessarily undesirable side effect of a more stringent, biologically relevant alignment yielding higher-quality counts more in sync with what was measured via sequencing. If one’s only metrics for evaluating an aligner are (i) the percentage of aligned reads and (ii) computational expense, then the ‘best’ aligner would be one which simply aligns every read to the first ncRNA in the reference database. Additionally, while it may be desirable to have a large proportion of reads align to reference ncRNAs, having this be the primary measure of an aligner’s utility underappreciates the inherent error rate in next-generation sequencing techniques [33].

Alignments to a genome and alignments to a database of reference ncRNAs are fundamentally different tasks and thus require different aligners. Bowtie, BWA, and STAR were designed to align reads to a genome where exact matches to reference sequences are not expected. In the different context of aligning reads from small RNA-sequenced data to a database of reference ncRNAs (where exact matches to entire reference sequences may be expected), we have demonstrated that the novel LevenMap tool yields more biologically relevant alignments than Bowtie, BWA, STAR, and BLAST.

## 5 Acknowledgements

## 5.1 Data and software availability

LevenMap is free, publicly available, and can be found on GitHub https://github.com/hdlugas/ LevenMap along with its documentation, implementation instructions, and a reproducible example. Feel free to issue a pull request if you have a suggestion for improvement or create an Issue if you encounter unexpected bugs. The publicly available data used in this study can be found in the European Nucleotide Archive with ID SRP335559 (https://www.ebi.ac.uk/ena/browser/view/SRP335559) [27].

## 5.2 Supplementary

The Supplementary File contains pseudocode for LevenMap along with tables and figures related to the differential expression analyses and the alignment characteristics of both the publicly available dataset and the prostate cancer cohort. The Supplementary file contains a table summarizing the computational expense of all aligners considered in this study as well.

## 5.3 Author contributions

H.D. and G.D. conceptualized the algorithm, H.D. developed the algorithm and performed the analysis, H.D., G.D., A.D, Y.K., and C.B. had discussions on issues common aligners such as Bowtie and BWA face when aligning small RNA-sequencing data to a database of ncRNAs, K.G. and J.B. were responsible for sample handling and RNA extraction and quantification, H.D. drafted the manuscript, C.B. was the PI of the DoD-funded project, and all authors reviewed and edited the manuscript.

