## Supplementary File for "Novel biologically relevant small RNA-sequencing alignment tool *LevenMap* for alignment to database of non-coding RNAs"

#### Contents

**1 Pseudocode** **1**
1.1 Build reference database . . . . . . . . . . . . . . . . . . . . . . . . 1
1.2 Alignment with LevenMap . . . . . . . . . . . . . . . . . . . . . . . 2

**2 Tables** **3**

**3 Figures** **5**

### 1 Pseudocode

#### 1.1 Build reference database

**Algorithm 1** Process reference database prior to alignment with LevenMap

**Input:**
FASTA file of reference ncRNAs
output_path: directory to which processed reference ncRNA database will be written
$N$: maximum number of mismatches allowed
$m$: minimum number of nucleotides
$M$: maximum number of nucleotides

**for** $i \leftarrow 0$ to $N$ **do**
  **for** $j \leftarrow m$ to $M$ **do**
    compute FASTA file of ncRNAs with length in interval $[j - i, j + 1]$
    processed FASTA file written to output_path
  **end for**
**end for**

#### 1.2 Alignment with LevenMap

**Algorithm 2** Alignment

```
Input:
  FASTQ file containing reads from small RNA-sequencing
  reference_database_path: path to processed reference ncRNA database
  N: maximum number of mismatches allowed
  output_file: tab-delimited text file of alignment output with one row for
               each aligned nucleotide sequence and three columns:
                   1. nucleotide sequence
                   2. reference sequence ID
                   3. number of reads corresponding to the nucleotide sequence

unique_reads ← all unique reads in the FASTQ file along with the number of duplicates

for i ← 1 to |unique_reads| do
    read ← unique_reads[i]
    reverse_complement_read ← reverse complement nucleotide sequence of read
    M ← number of nucleotides in read
    references ← all reference ncRNAs with length in [M − N, M + N]
    min_dist = ∞

    aligned_refs ← “”
    for ref_seq in references do
        ref_ID ← ID corresponding to ref_seq
        dist_forward ← Levenshtein_distance(read,ref_seq)
        dist_reverse_complement ← Levenshtein_distance(reverse_complement_read,ref_seq)
        min_d ← min(dist_forward,dist_reverse_complement)

        if min_d < min_dist then
            aligned_refs ← “”
            min_dist ← min_d
        else if min_d == min_dist then
            aligned_refs ← concatenate(aligned_refs,‘&’,ref_ID)
        end if

        if min_dist ≤ N then
            N_duplicates ← number of duplicates of read in FASTQ file
            row ← [read, ref_ID, N_duplicates]
            append row to output_file
        end if
    end for
end for
```

#### 2 Tables

**Table S1:** Prostate cancer cohort patient characteristics.

| Variable | Level | Value (N=34) |
|---|---|---|
| Age at Biopsy | | 64 $\pm$ 7 |
| Race | Black | 9 (26.5%) |
| | White | 25 (73.5%) |
| Targeted Site PI-RADS Score | 3 | 6 (20.0%) |
| | 4 | 12 (40.0%) |
| | 5 | 12 (40.0%) |
| Targeted Site Grade | 0 | 16 (47.1%) |
| | 1 | 9 (26.5%) |
| | 2 | 3 (8.8%) |
| | 3 | 4 (11.8%) |
| | 4 | 1 (2.9%) |
| | 5 | 1 (2.9%) |
| Cancer Status | Cancer | 18 (52.9%) |
| | Not Cancer | 16 (47.1%) |

**Table S2:** Computational expense of each aligner applied to (i) the in-house prostate cancer cohort and (ii) the publicly available SRP335559 dataset. The computation times reported are mean ± standard deviation across all samples with units of minutes. Non-LevenMap aligners were run using 1 CPU core on an x86_64 AMD system. Flag: indicates whether or not to suppress alignments if multiple alignments exist (T: only alignments of reads with exactly one alignment are reported; F: all alignments are reported). L: seed length.

| Aligner | Prostate Cancer Cohort | SRP335559 |
|---|---|---|
| LevenMap: no mismatches, 1 CPU | 20.243 ± 16.548 | 21.730 ± 11.159 |
| LevenMap: at most 1 mismatch, 1 CPU | 37.676 ± 22.837 | 59.028 ± 37.465 |
| LevenMap: at most 2 mismatches, 1 CPU | 57.783 ± 24.455 | 81.342 ± 50.699 |
| LevenMap: no mismatches, 4 CPUs | 2.376 ± 0.868 | 6.448 ± 4.516 |
| LevenMap: at most 1 mismatch, 4 CPUs | 6.207 ± 2.411 | 16.807 ± 13.116 |
| LevenMap: at most 2 mismatches, 4 CPUs | 10.060 ± 3.944 | 25.973 ± 21.257 |
| Bowtie: Flag=T, L=28 | 0.394 ± 0.187 | 1.587 ± 0.706 |
| Bowtie: Flag=T, L=10 | 0.587 ± 0.292 | 1.945 ± 0.805 |
| Bowtie: Flag=F, L=28 | 0.327 ± 0.363 | 1.173 ± 0.495 |
| Bowtie: Flag=F, L=10 | 0.523 ± 0.256 | 1.646 ± 0.684 |
| BWA: L=32 | 5.622 ± 3.336 | 7.988 ± 3.078 |
| BWA: L=10 | 5.737 ± 3.396 | 8.112 ± 3.274 |
| STAR | 24.003 ± 11.056 | 42.158 ± 15.780 |
| BLAST | 151.272 ± 246.552 | 268.026 ± 529.134 |

### 3 Figures

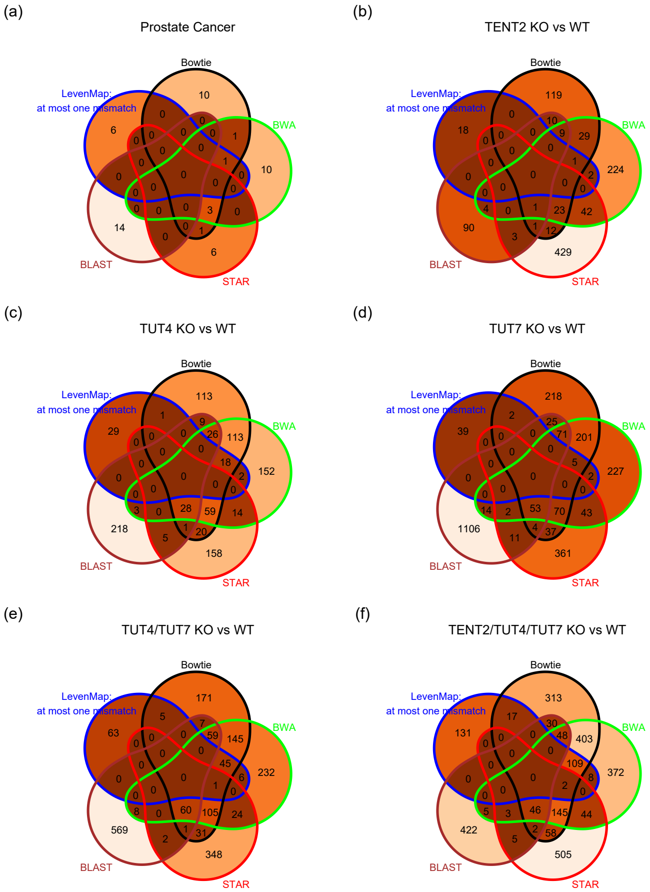


**Figure S1:** Venn diagrams depicting the commonality of differentially expressed ncRNAs among all aligners. **(a)**: Prostate cancer cohort targeted site cancer status. **(b)-(f)**: SRP335559 dataset TENT2 knock-out vs wild-type, TUT4 knock-out vs wild-type, TUT7 knock-out vs wild-type, TUT4/TUT7 knock-out vs wild-type, and TENT2/TUT4/TUT7 knock-out vs wild-type, respectively.

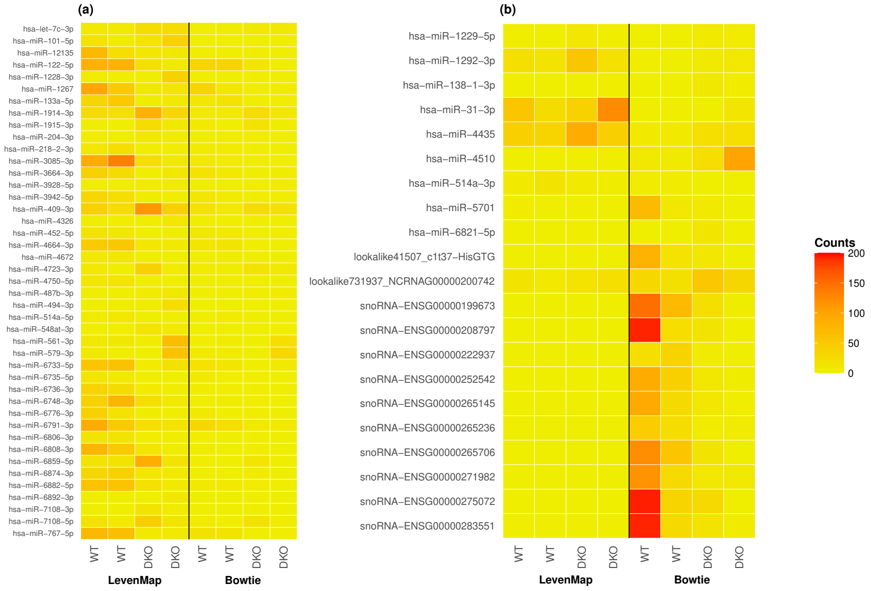


**Figure S2:** Heatmaps depicting the raw counts of ncRNAs with respect to both LevenMap (with at most one mismatch allowed) and Bowtie (with all alignments reported and a seed length of 10) in the SRP335559 dataset WT and DKO samples. **(a)**: ncRNAs differentially expressed (i.e. those with nominal p-value less than 0.05 and $|\log_2(\text{fold-change}) > 1|$) with respect to LevenMap and not Bowtie are shown. **(b)**: ncRNAs differentially expressed with respect to Bowtie and not LevenMap. Additionally, only ncRNAs with a total count across all samples in a given comparison less than 300 are shown to exclude outliers which render differentially expressed ncRNAs with fewer overall counts difficult to visually examine in the figure. WT: wild-type. DKO: double KO (TUT4/TUT7 KO).

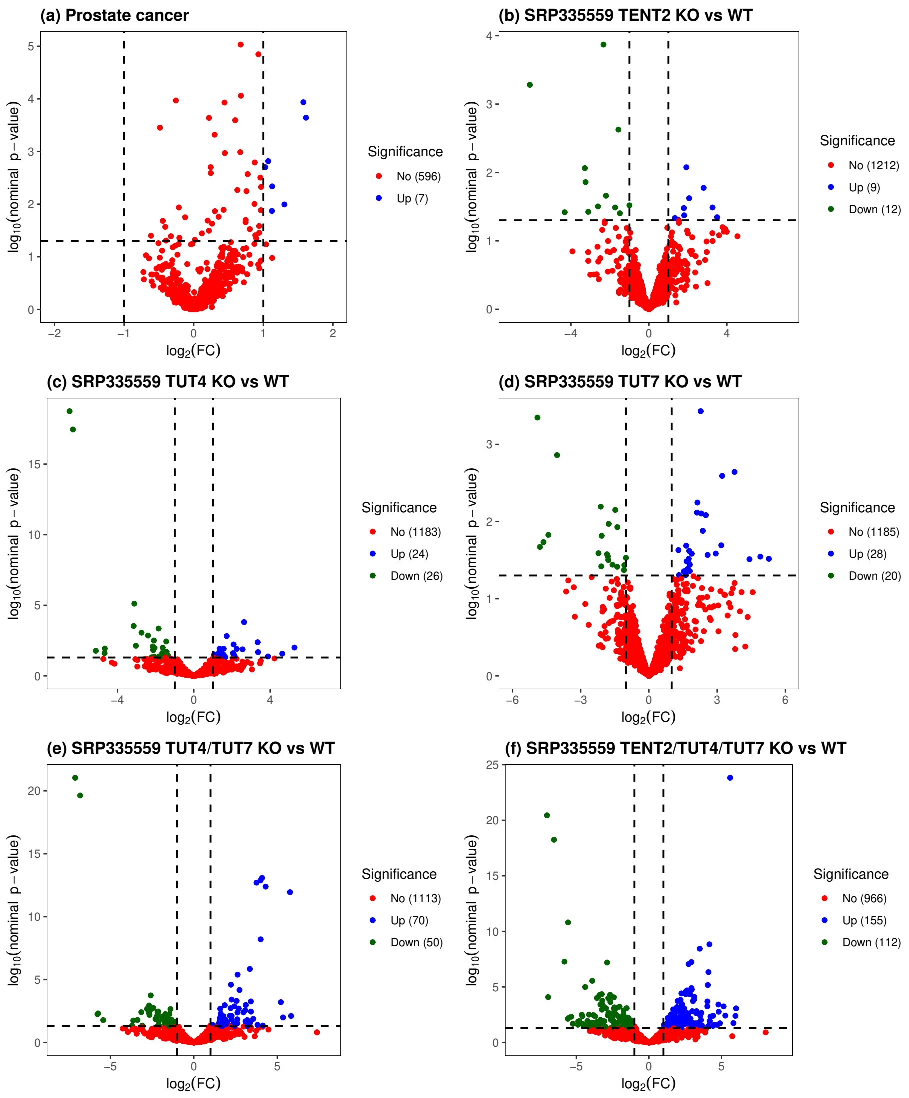


**Figure S3:** Volcano plots with respect to the LevenMap alignment allowing for at most one mismatch. Each point represents a ncRNA. **(a)**: Results from the generalized least squares differential expression analysis of the prostate cancer cohort. **(b)-(f)**: Results from the DESeq2 differential expression analysis of the SRP335559 dataset comparing TENT2 knock-out vs wild-type, TUT4 knock-out vs wild-type, TUT7 knock-out vs wild-type, TUT4/TUT7 knock-out vs wild-type, and TENT2/TUT4/TUT7 knock-out vs wild-type, respectively.
